# Machine Learning-Guided Classification of Druggable Pockets and Phylogenetic Druggability Transfer Across the Human Kinome

**DOI:** 10.64898/2026.09.06.749698

**Authors:** Rudrakshi Chauhan, Rajiah Aldrin Denny, N. Arul Murugan

## Abstract

Protein kinases are among the most intensively pursued therapeutic targets in oncology and beyond, yet selectivity remains largely unsolved; around 536 Manning kinase domains share a conserved ATP-binding pocket, making it difficult to target one without hitting others. Allosteric binding modes which exploit conformational states unique to individual kinases or narrow kinase subfamilies offer a principled route to selectivity. Yet the absence of a kinome-wide structural landscape of these pockets limits their translational potential. Here we construct a machine learning-guided structural atlas of 11,945 kinase-inhibitor complexes, training an Extra Trees classifier on pocket residue interaction with ligand to assign all seven canonical binding modes and resolve allosteric subclasses with pharmacological precision. Our structural analysis reveals that approximately 303 Manning kinase domains have known inhibitors bound to them, representing ∼56.5% of the Manning kinase domain; of these, only 26% (78 kinases) are targeted by non-ATP-competitive allosteric inhibitors, indicating substantial unexplored pharmacological space. Integrating these classifications with the Manning kinome phylogeny, we demonstrate that evolutionary proximity is a statistically significant predictor of shared allosteric pocket accessibility. Because a functionally similar target has similar conformational dynamics and similar druggable pockets, ‘phylogenetic druggability transfer’ can act as a strong signal for identifying whether a given kinase can be targeted using an allosteric inhibitor or not. In addition, this work repositions the kinome phylogeny as a map of pharmacological opportunity and provides a reusable predictive framework for drug discovery programs - including orthosteric, allosteric, covalent, bifunctional inhibitors, and chemical degrader - across the understudied kinome.

**Significance Statement:** Most kinase drugs bind to pocket shared across the kinome, making selectivity difficult. Allosteric pockets are more selective, but their presence and accessibility are not understood across kinome. Testing a candidate against a specific binding mode requires a dedicated assay, which is expensive and time consuming, and wasted effort if that pocket is not actually accessible. We classified thousands of kinase structures by binding mode and mapped them onto the kinome phylogenetic tree. Closely related kinases often share the same druggable pockets, so this map allows us to predict likely pockets in related kinases, helping prioritize assays and identify repurposing opportunities, making drug discovery more efficient.

## 1. Introduction

Protein kinases are essential regulators of cellular signaling pathways, catalyzing the phosphorylation of substrate proteins by transferring the γ-phosphate of ATP, thereby regulating protein activity, localization, and interaction networks across virtually every cellular signaling cascade[1], [2]. Through this post-translational modification, kinases orchestrate cell proliferation, differentiation, survival, metabolism, and apoptosis[3], [4]. The approximately 518 human protein kinases are organized into eight canonical phylogenetic groups (AGC, CAMK, CK1, CMGC, RGC, STE, TK, TKL) plus atypical kinases, as defined by Manning et al. based on sequence and structural similarity[5]. Tight regulation of kinase activity is essential due to their central function in cellular homeostasis, dysregulation of kinases via somatic mutation, overexpression, or truncation can drive pathological signaling underlying cancer, autoimmune disease, and neurodegeneration[1], [4], [6]. Over the past two decades, protein kinases have emerged as the most productive and clinically impactful drug target class in modern pharmacology, with kinase accounting for 62 of 802 successful therapeutic targets[7] with more than >130 drugs approved worldwide[8]. The approval of imatinib in 2001, exploiting the unique DFG-out ATP non-competitive conformation of BCR-ABL, transformed the treatment of chronic myeloid leukemia and established the paradigm of conformation-selective kinase targeting[9]. However, the vast majority of approved drugs are ATP-competitive (Type I) inhibitors that suffer from limited selectivity owing to high conservation of the ATP-binding hinge interface across kinase families[6], [10]. This promiscuity drives off-target toxicity, contributes to acquired drug resistance, and creates a persistent bottleneck in precision oncology that the field is actively resolving through allosteric and induced-proximity approaches[8], [11].

Induced-proximity pharmacology, particularly molecular glue and protein degraders, represent an emerging strategy in kinase drug discovery, where small molecules induce protein-protein interactions that redirect E3 ubiquitin ligases to target previously inaccessible proteins for proteasomal degradation. The clearest example within the kinase field is CR8, a CDK inhibitor whose kinase bound form exposes a surface moiety that bridges CDK12-cyclin K directly to the CUL4 adaptor protein DDB1, triggering cyclin K degradation without requiring a canonical DCAF substrate receptor[12]. Subsequent work showed that other CDK12 inhibitors, including SR-4835, operate through the same principle via their surface-exposed dichlorobenzimidazole side chain[13]. Importantly, all characterized CDK12 molecular glues to date engage exclusively the ATP binding pocket, what our classification defines as Type I binding modes. This means that the surfaces available to recruit degradation machinery are limited to those accessible from the front pocket. The binding mode atlas and phylogenetic druggability framework developed here suggest that allosteric pockets could anchor molecular glue scaffolds that project chemically distinct surfaces, offering routes to induced proximity degraders with improved selectivity for kinases where ATP site based approaches remain limited.

A central insight driving this work is that kinase function is inseparable from structure and conformational dynamics. Kinases do not exist in a single static conformation but sample an ensemble of conformational states that differ in catalytic competence, regulatory interactions, and crucially in the binding pockets they expose[14], [15]. The hydrophobic regulatory spine, four non contiguous hydrophobic residues assembled in the active conformation, identified by Kornev et al. as the structural hallmark of kinase activation[16], [17], is the architectural determinant of conformational switching: DFG-Phe occupies the spine in the active state, and its rotation upon DFG flip creates the cryptic allosteric back pocket exploited by Type II inhibitors[17], [18]. These conformational degrees of freedom are not randomly distributed across the kinome but are shaped by evolutionary history. The structural plasticity of a given kinase, its propensity to sample specific inactive conformations and expose specific allosteric pockets, is a consequence of its catalytic domain sequence, which is phylogenetically determined[19], [20]. This convergence of structural biology and evolutionary analysis creates a powerful logical framework. If pocket accessibility is a phylogenetically conserved structural property, then experimental evidence of pocket accessibility in one kinase predicts the likely pocket accessibility of its evolutionary relatives which is known as phylogenetic druggability transfer. It has been implicitly exploited historically, the development of imatinib analogs against the phylogenetically related ABL, KIT, and PDGFR kinases is the clearest example[9]. But this has never been formalized or tested systematically at the full kinome scale across all inhibitor binding mode classes. Beyond structural conservation, phylogenetically related kinases also share functional properties: they participate in overlapping signaling pathways, phosphorylate related substrate motifs[19], [20], and exhibit similar patterns of oncogenic mutation and disease association[21]. This means that pharmacological exploitation of a shared allosteric pocket in phylogenetically related kinases may simultaneously address pathway level functional redundancy, a major driver of acquired resistance to targeted therapy[22], [23].

Existing computational approaches to kinase binding mode classification have several limitations that prevent full realization of this potential. Rule-based classification schemes, as implemented in the structural annotation of pockets in KLIFS[24], [25], relies on a fixed set of geometric criteria that cannot capture the full structural diversity of kinase-ligand interactions, leaving approximately 24% of entries unclassified. Similarly, KinCore[26] uses a rule based scheme in which ligands that do not satisfy the criteria for any defined class are assigned to Type I by default that is a conservative fallback that ensures every entry receives a label, but one that can obscure cases where the true binding mode differs from Type I. To characterize the practical impact of this design choice, we compared classifications between our method and KinCore across a shared set of ligand protein pairs (11,264). Of these, 10,480 (93.0%) were classified identically by both approaches, indicating strong overall agreement. Among the 784 mismatches, 646 corresponded to disagreements between Type I and Type I.5 assignments - two classes whose structural distinction is inherently subtle and often differentiated by relatively minor conformational details. These cases reflect the difficulty of drawing a hard boundary between closely related binding modes rather than a substantive disagreement between methods. The remaining 138 mismatches, however, represent more meaningful discrepancies, of these, approximately 100 corresponded to cases where KinCore’s default of Type I fallback assigned a label in the absence of a clear rule match, whereas our method identified a distinct binding mode based on the underlying structural features, suggesting that while rule based frameworks such as KLIFS and KinCore have been instrumental in establishing kinase binding mode taxonomies, their reliance on fixed decision rules and default fallbacks leaves a meaningful fraction ambiguously classified.

Prior machine learning approaches, including Miljković et al. [27] and Abdelbaky et al. [28]. were limited to four binding mode categories using ligand only feature sets, leaving the multi class allosteric subclass landscape unmapped. No prior study has integrated comprehensive ML based binding mode classification with kinome phylogeny to systematically test and operationalize the phylogenetic druggability transfer hypothesis.

Here, we address these gaps through four principal contributions: (i) A comprehensive organism-spanning kinase structural atlas assembled for systematic inhibitor classification (11,945 ligand protein pair, and structures from 477 kinase domain UniPro); (ii) a high accuracy classifier model covering all five binding mode classes and three allosteric subclasses; (iii) pharmacological validation of all allosteric pocket subclasses through approved or clinically advanced inhibitor structures; and (iv) full kinome statistical validation of the phylogenetic druggability transfer principle, with quantitative demonstration that evolutionarily related kinases share not only pocket geometry but also functional conformational accessibility. The drug repurposing framework of Chakraborti et al. [29] demonstrating that evolutionary proximity between drug targets enables systematic drug repurposing, and the contemporary review of kinase modulation strategies by Dar et al. [8] identifying allosteric and induced proximity approaches as the frontier of kinase drug discovery, together provide the conceptual scaffolding for this work.

### 1.1. Kinase Structural Features and Inhibitor Classification

Despite their functional diversity, protein kinases share a highly conserved bi-lobal catalytic domain with a smaller N terminal lobe (N-lobe) rich in β-sheets and a larger C terminal lobe (C-lobe) dominated by α-helices, connected by a hinge region that forms part of the ATP binding cleft[10]. Within this conserved scaffold, the kinase fold supports remarkable conformational diversity through the coordinated movement of several key regulatory structural elements.

The glycine rich P-loop (typically GxGxxG) stabilizes the β and γ-phosphates of ATP and exhibits conformational flexibility accommodating diverse ATP competitive ligands[10]. The regulatory and catalytic hydrophobic spines, identified by Kornev et al. as dynamically assembled sets of non-contiguous hydrophobic residues, provide the architectural basis for kinase activation and define the intramolecular communication network transmitting allosteric signals from distal regulatory sites to the active site[16], [17]. Disruption of the regulatory spine is the structural mechanism underlying DFG-out pocket formation. Phe residue in DFG occupies the spine position in the active state, and its rotation upon DFG flip exposes the cryptic allosteric back pocket[17], [18]. The αC-helix, containing the conserved glutamate that forms the catalytic salt bridge with the β-III lysine, serves as a signal integration motif[17]. The activation loop phosphorylation state regulates substrate access, while the DFG (Asp-Phe-Gly) motif coordinates a magnesium ion essential for catalysis[14], [18].

Critically, the propensity to sample specific conformational states is not uniform across the kinome but is phylogenetically stratified, because the residues determining conformational accessibility, including the x-DFG position, the gatekeeper residue, the hydrophobic spine composition, and regulatory domain geometry, are evolutionarily conserved within kinase clades and diverge between distant groups[19], [30]. This means that structural plasticity and the resulting pocket accessibility are heritable properties of kinase clades, a principle at the foundation of the phylogenetic druggability transfer framework.

The five class kinase inhibitor classification framework, provides the pharmacological vocabulary for this work. Type I inhibitors (DFG-in, αC-in, ATP-competitive) are broadly distributed but exhibit limited selectivit[10], [31]. Type I½ inhibitors (DFG-in, αC-out, extended back-pocket binding) offer improved selectivity through αC-helix displacement[10], [11]. Type II inhibitors (DFG-out, back-pocket engagement) leverage the structural uniqueness of the inactive conformation[8], [10]. Type III inhibitors (allosteric, ATP-non-competitive) achieve exceptional selectivity through targeting less conserved adjacent allosteric pockets[10], [11]. Type IV inhibitors (remote allosteric) engage distal regulatory sites, exemplified by asciminib’s clinical validation as the first specifically allosteric kinase inhibitor[8], [32]. As Rudolph et al. document in their 2026 review, allosteric and induced proximity approaches are now the dominant frontier of kinase drug discovery precisely because they circumvent the selectivity bottleneck of ATP site conservation[8].

## 2. Methodology

The workflow proceeds through four main stages (i) systematic retrieval and curation of kinase domain structures from primary databases, (ii) standardized computation of structural descriptors capturing binding pocket geometry and conformational states of the kinase (iii) Rule based annotation of ligand binding modes, followed by manual structural visualization and inspection to verify conformational assignments, yielding a high confidence labeled training dataset and (iv) development of a supervised machine learning classifier to extend classification coverage to all curated kinase ligand complexes, followed by integration with kinome phylogeny for druggability transfer analysis.

### 2.1. Dataset Construction and Curation

To establish a comprehensive and organism spanning foundation for the structural atlas, all reviewed (Swiss-Prot) UniProt entries were systematically retrieved from the UniProt database[33]. For each UniProt accession, domain topology annotations were extracted and entries containing at least one kinase-related domain were retained. Specifically, domains matching the annotations "protein kinase,"and "PI3K/PI4K catalytic domain" were selected, consistent with established domain nomenclature for the eukaryotic protein kinase (ePK) superfamily and atypical kinases (aPKs)[5]. The residue boundaries defining each annotated kinase domain were recorded to enable downstream domain level structural mapping. Importantly, a single UniProt entry could contribute multiple kinase domain records, ensuring that multi domain kinase proteins were handled at the individual domain level throughout all analyses. For each UniProt accession, all deposited Protein Data Bank (PDB) structures were identified through the UniProt PDB cross reference mapping only [33], [34]. Only PDB chains whose resolved residue sequences overlapped with the annotated kinase domain boundaries were retained. This initial query yielded 17,490 PDB chain instances associated with 692 UniProt accessions and 713 UniProt kinase domain entries. To ensure that structures contained sufficient coverage of the kinase domain for meaningful structural analysis, a domain coverage threshold of ≥30% was applied, retaining 12,923 PDB chain instances corresponding to 8,548 unique PDB entries, mapped across 479 UniProt accessions. Each PDB chain was treated as an independent structural unit throughout the pipeline. In cases where a single PDB entry contributed multiple chains, each chain was processed separately, consistent with the possibility of distinct crystal packing environments, ligand occupancy states, or conformational states between chains within the same deposited entry. Similarly, chains annotated with more than one kinase domain were handled at the individual domain level.

To enable consistent cross kinase structural comparison, the established 85 residue binding site framework from the KLIFS database was employed[24], [25]. Multiple sequence alignment was performed across all UniProt kinase domain sequences using a structure informed alignment approach, guided by conserved catalytic motifs (DFG, HRD, APE) and secondary structural elements common to the kinase fold[35] and shared secondary structural elements of the kinase fold. Based on this alignment, the predefined set of 85 topologically conserved residue positions representing the ATP-binding pocket and its immediate flanking regions was utilized (Figure 1). These positions encompass the key functional sub regions of the catalytic cleft, including the glycine-rich P-loop (positions 4-9), αC-helix (positions 20-30), back loop (positions 31-37), gatekeeper residue (position 45), hinge region (positions 46-48), linker (positions 49-52), catalytic loop (positions 68-75), DFG motif and its upstream xDFG position (positions 80-83), and activation loop (positions 84-85) (Figure 1). For each structure, the 85 conserved binding site residues were mapped onto the PDB chain coordinates using UniProt based residue indexing. Structures in which fewer than 30% of the 85 defined binding site residues were resolved and mappable were excluded from further analysis. This secondary filtering step yielded a final curated dataset of 12,641 kinase domain structures with high quality, consistently mappable binding site coverage.

**Figure 1.**
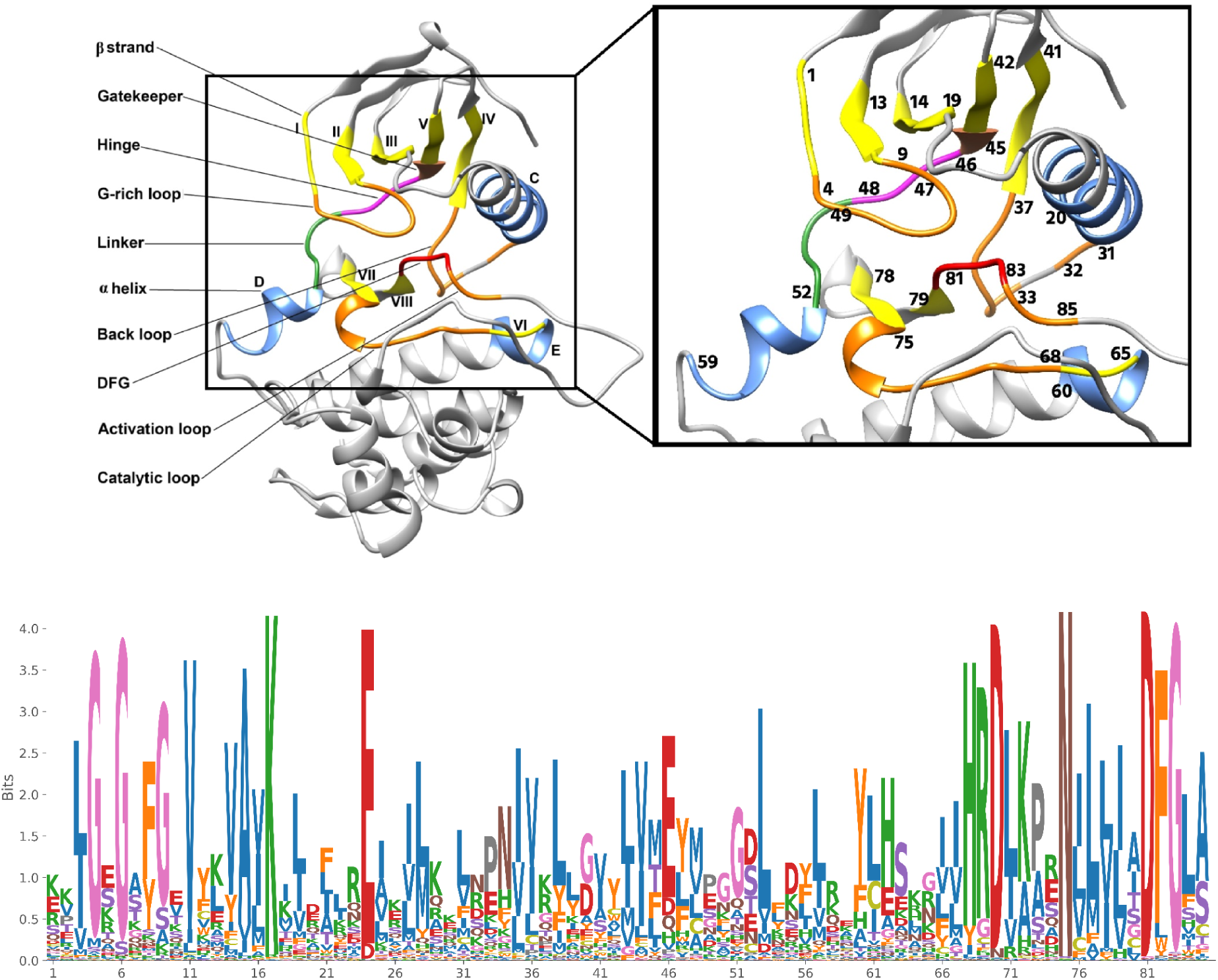
The 85 residue KLIFS binding site framework mapped onto the kinase catalytic domain. (Left) Ribbon representation of the kinase domain with sub-regions colour-coded: hinge (magenta), helix (blue), sheets (yellow) and DFG motif (red). (Right) Zoomed view with individual position numbers annotated. (Bottom) sequence logo of conservation across all UniProt kinase domains at each of the 85 positions.

To focus the analysis classification of kinase inhibitor complexes, the curated structural dataset was filtered to retain only structures containing at least one bound small molecule ligand. Common solvent molecules, buffer components, crystallographic additives and metal ions were systematically excluded. Ligand identity was assigned based on the PDB Chemical Component Dictionary[34]. This filtering step identified 10,739 structures containing at least one qualifying ligand. Because a single PDB structure or chain may contain multiple bound ligands, each unique ligand molecule within a structure was treated as an independent data point. Structures were therefore decomposed into individual protein-ligand pair records where applicable. This procedure yielded a final dataset of 11,945 unique kinase-ligand complex records, each defined by a specific PDB entry, chain identifier, and ligand instance. This dataset served as the primary analytical unit for all descriptor calculation, binding mode classification, and machine learning analyses.

### 2.2. Structural Descriptor Calculation

For each kinase ligand complex, a quantitative representation of the spatial relationship between the ligand and the 85 residue binding site framework was computed. Euclidean distances were calculated between all heavy atoms of the ligand molecule and the heavy atoms of each of the 85 mapped binding site residues. For each residue position, the minimum heavy-atom distance to any ligand atom was retained as the descriptor value, providing a continuous, residue-level measure of ligand proximity that preserves the geometric arrangement of the interaction without imposing arbitrary contact cutoffs. This procedure produced an 85-dimensional continuous distance feature vector for each kinase-ligand complex, enabling finer discrimination of binding geometry and interaction strength. These distance vectors served as the primary feature input for the classification.

The conformational state of the DFG (Asp-Phe-Gly) motif; a primary determinant of kinase activation state and the key structural switch governing access to the Type II allosteric back pocket[36], was characterized using a local geometric coordinate system defined for each structure. The origin of this coordinate frame was placed at the Cα atom of the terminal residue of β-strand VIII, immediately preceding the DFG motif (VIII.79). The principal x-axis was defined by the unit vector from VIII.79 to Aspartate residue of the DFG Motif (xDFG.80), and a local reference plane was constructed using the Cα atom of the conserved αC-helix residue Glutamate (αC.24) as an out-of-plane reference point. Using this coordinate frame, displacement vectors from the origin to each DFG residue (xDFG.81/Asp, xDFG.82/Phe, xDFG.83/Gly) were computed, along with their magnitudes and pairwise inter-residue angles. Additional geometric descriptors capturing the overall torsional state of the DFG motif were included as indicators of DFG flip geometry. Together, these descriptors enabled robust classification of each structure into one of three DFG conformational states: DFG-in, DFG-out, or DFG-out-like (an intermediate conformation observed in structures where the phenylalanine side chain has begun to rotate but has not completed the full DFG flip)[14].

The position of the αC-helix relative to the active site was quantified by measuring the Cα-Cα distance between the conserved catalytic lysine in β-strand III (III.17) and the conserved glutamate in the αC-helix (αC.24). This inter-residue distance serves as a proxy for αC-helix displacement[37]. Structures with an III.17-αC.24 Cα distance of ≤13.15 Å were assigned to the αC-in state (active-like, permitting salt bridge formation), while those exceeding this threshold were assigned to the αC-out state (inactive-like, salt bridge disrupted). To directly assess salt bridge formation between the catalytic lysine and αC-helix glutamate, the distances between the NZ atom of III.17 and the carboxylate oxygen atoms (OE1 and OE2) of αC.24 were measured. A salt bridge was considered present if either NZ-OE distance was ≤4.0 Å, consistent with standard criteria for electrostatic contact identification in structural biology[38].

Each kinase structure was assigned an overall activation state - Active, Intermediate, or Inactive, based on the combined conformational descriptors of the DFG motif, αC-helix position, and salt bridge status, following a hierarchical decision logic. A structure was classified as Active when all three criteria were simultaneously satisfied that are formation of the salt bridge between the β-sheet III lysine and αC-helix glutamate, adoption of a DFG-in or conformation, and positioning of the αC-helix in the "in" state. Structures capturing partially-activated configurations with absent salt bridge or DFG-out-like geometry while the αC-helix retained its "in" position or when the αC-helix is in "out" position but the DFG motif adopted a DFG-in geometry. Structures were classified as Inactive when the DFG motif adopted a DFG-out conformation, or when αC-helix "out" coincided with a DFG-out-like geometry (Table S1). This framework captures the combinatorial nature of kinase regulation, recognizing that neither the DFG conformation nor the αC-helix position alone is sufficient to define activation state. Both structural elements must be considered jointly, as has been established by systematic studies of the kinase conformational landscape[14], [39].

**Figure 2.**
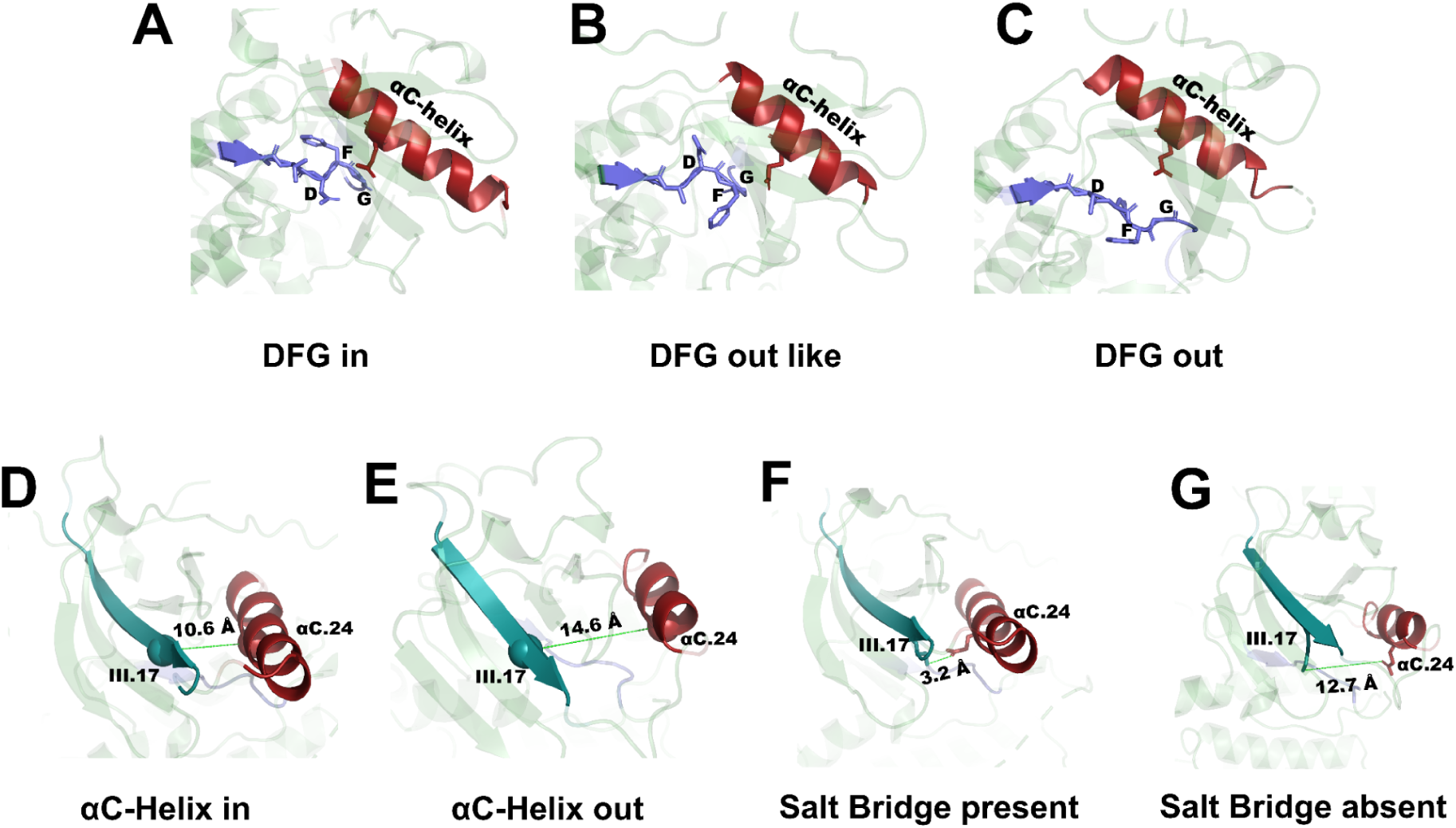
Structural determinants used for kinase activation-state annotation. Representative DFG-in (A), DFG-out-like (B) and DFG-out (C) conformations depicted as ribbon structures with the DFG tripeptide shown as sticks. αC-helix-in (D) and αC-helix-out (E) states. The catalytic salt bridge between β-strand III lysine (III.17) and αC-helix glutamate (αC.24) Presence (F) and absence (G). with distances shown as dashed lines.

### 2.3 Rule-Based Annotation of Inhibitor Binding Modes

To generate a high confidence labeled dataset for supervised machine learning, a deterministic, rule-based classification scheme was used, grounded in established mechanistic principles of kinase inhibitor pharmacology[10], [40]. Each kinase-ligand complex was evaluated against a hierarchically organized set of structural criteria encompassing the spatial occupancy of the ligand across defined sub-regions of the binding pocket (front pocket, gate area, back pocket, and distal allosteric regions) based on the KLIFS pocket partitioning scheme[25], The conformational state of the DFG motif and αC-helix, and the presence of specific interaction types, such as hinge-region hydrogen bonds.

#### Type I Inhibitors

Type I inhibitors were defined as ligands occupying the front subdivision of the ATP-binding pocket, with no extension into the back pocket. Required criteria: (i) DFG conformation in the DFG-in or DFG-out-like state, (ii) αC-helix in the αC-in conformation, and (iii) at least 1 bond contact with the hinge residues, reflecting canonical ATP-competitive binding. This category encompasses the most common class of approved kinase inhibitors and represents the baseline binding mode for ATP-competitive agents(figure 3A).

**Figure 3.**
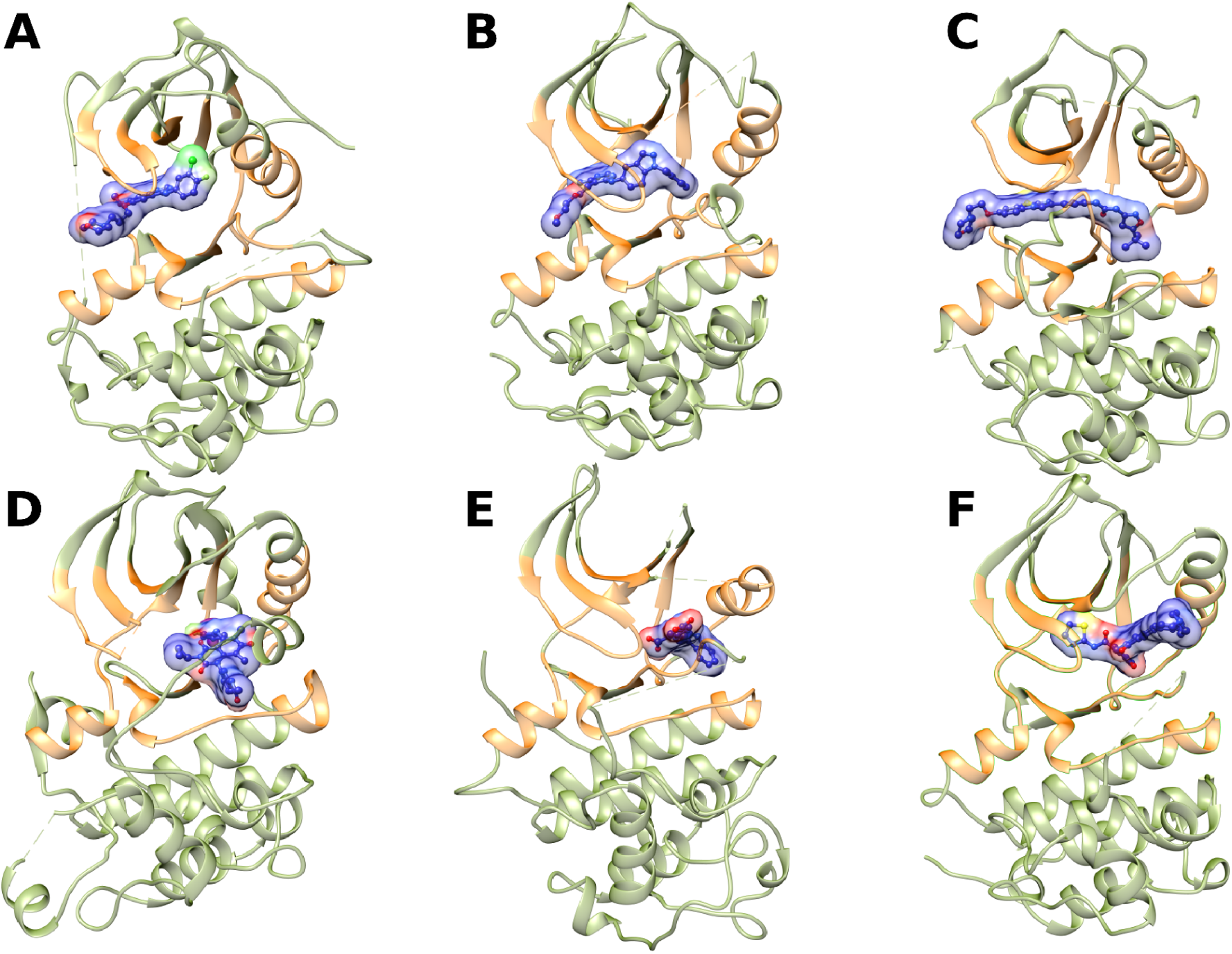
Representative structures of the five kinase inhibitor binding-mode classes. Co-crystal structures showing Type I (**2ITO**) (A), Type I½ (**3BEL**) (B), Type II (**4XUF**) (C), Type III-Top (**7JUX**) (D), Type III-Back (**8FP0**) (E) and Type III-Front (**7LG8**) (F) inhibitors (blue surface) bound to the kinase catalytic domain (orange). Each panel illustrates the characteristic pocket occupancy and conformational state defining each class.

#### Type I½ Inhibitors

Type I½ inhibitors were defined as ligands bridging the front pocket and gate area, with limited, permitted extension into the back pocket, region bounded by the DFG motif, β-strands III and V, the β-III conserved lysine, and the αC-helix. Required criteria: (i) DFG-in conformation, (ii) αC-helix in the αC-out conformation, reflecting partial displacement, and (iii) at least 1 bond contact with the hinge residues(figure 3B).

#### Type II Inhibitors

Type II inhibitors were defined as ligands occupying the front pocket, gate area, and back pocket simultaneously, requiring interaction with the allosteric region between the DFG motif and αC-helix. Required criteria: (i) DFG-out conformation, (ii) αC-helix in either αC-in or αC-out state, and (iii) at least 1 bond contact with the hinge residues. The DFG-out conformation is non-negotiable for this class, as it is the conformational prerequisite for back pocket exposure (figure 3C).

#### Type III Inhibitors

Type III inhibitors were defined as allosteric ligands occupying regions adjacent to the ATP-binding site without ATP-competitive binding. Required criteria: (i) no occupancy of the canonical ATP front pocket, (ii) exclusive or predominant interaction with the back pocket, and (iii) absence of hinge region hydrogen bonds. To capture the spatial diversity of Type III binding modes, this class was further resolved into three spatially defined subclasses based on the ligand’s positional relationship with the activation loop: Type III-Top where the ligands is positioned above the activation loop within the catalytic pocket, bridging the active site and adjacent allosteric space (figure 3D). These compounds may serve as structural templates for Type II inhibitor design. Type III-Back are ligands positioned entirely behind the activation loop, fully occupying the back pocket without any extension toward the front pocket. These represent the most purely allosteric configurations within this class (figure 3E). And Type III-Front, In this subclass ligands occupy the back pocket with extension over the activation loop toward the front region of the binding site, spanning the loop boundary while retaining non-ATP-competitive character (figure 3F).

#### Type IV Inhibitors

Type IV inhibitors were defined as ligands binding entirely outside the ATP-binding cleft and all defined subdivisions of the catalytic pocket, engaging distal allosteric sites within or at the surface of the kinase domain. Required criterion for this class used were: no spatial overlap with any sub-region of the 85-residue binding pocket framework. To capture the structural diversity of remote allosteric binding modes, Type IV inhibitors can be further resolved into six major spatially distinct subclasses based on their binding locations within discrete regions of the kinase domain (figure 7 A-F).

Any kinase-ligand complex that did not satisfy the complete set of required criteria for any of the defined binding mode categories was designated as Unclassified. This conservative approach prioritized classification specificity over sensitivity, ensuring that all rule-based annotations carried high structural confidence. This Unclassified data isn’t noise or failure but actually the ambiguous cases that become the prediction target for the ML model and were excluded from the supervised machine learning training set, while the high-confidence rule-based labels become the training data (ground truth) to train that ML model on.

### 2.4 Limitations of Rule-Based Classification

The deterministic rules relied on rigid distance cutoffs and binary conformational labels. This division of continuous structural features is problematic for kinase-ligand systems, which frequently display intermediate or ambiguous states. For example, hinge interactions were only recognized if the ligand-residue distance was <4.0 Å, thereby excluding weaker but still stabilizing contacts. Similarly, αC-helix was forced into in vs. out categories, overlooking ambiguous intermediate or partially shifted conformations in the PDB. Rules also depended on a small set of residues (e.g., DFG, αC-helix and hinge.), so missing density or minor coordinate errors were sufficient to break classification. Finally, the rule-based scheme gave only hard labels, with no way to handle uncertainty. Complexes that fell in borderline cases were simply discarded as unclassified. Complexes such as 1yxu and 3ii5 lacked recognized hinge contacts yet are Type I/II inhibitors, whereas 5r91 does have hinge contacts but the inhibitor is type 4. Missing structural elements also caused failures like in pdb 2qo7, hinge residues were unresolved, while in 8tze and 5owq, the DFG motif was absent. In other cases, conformational descriptors contradicted experimental evidence. For example, 1b38 and 3rhk were rejected because αC-helix was “in” and DFG was “out” for 1b38 and both the DFG and αC-helix in 3rhk were labeled “out”, despite being Type I inhibitors. Similarly, 5tco and 3b2w are Type II inhibitors but were assigned as unclassified due to “in” annotations of both the DFG and αC-helix.

To illustrate how machine learning resolves the limitations described, we examined specific complexes that were unclassified by deterministic rules. The following figure 4, summarizes representative examples.

**Figure 4.**
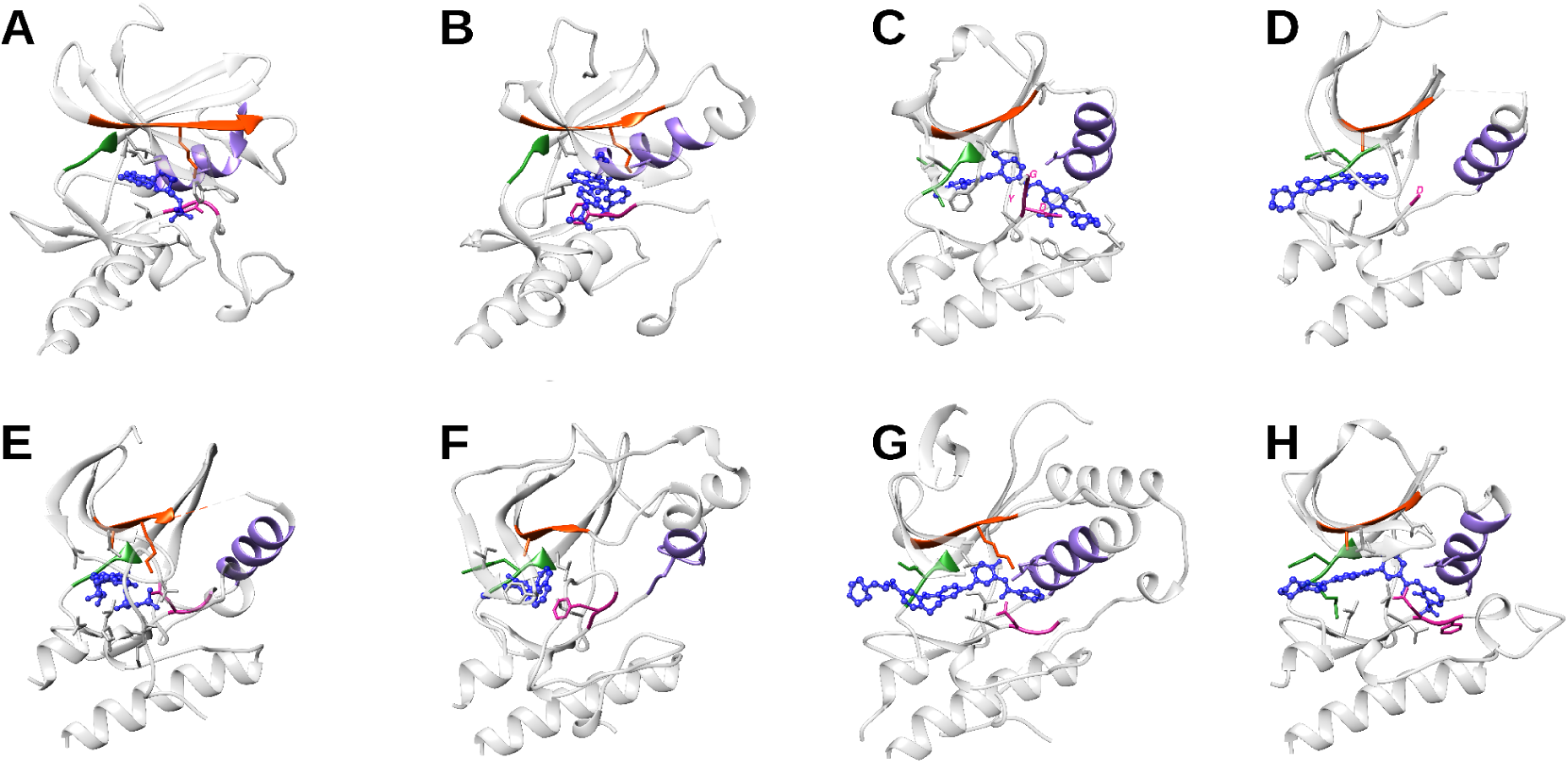
Representative kinase-inhibitor complexes highlighting the limitations of rule-based classification where the complexes were left Unclassified. In panels A (**1YXU**) and B (**3II5**), inhibitors are type I and type II, respectively, but both lack hinge contacts, resulting in an unclassified label. In panel C (**8TZE**), a point mutation causes misannotation of the DFG motif, while in panel D (**5OWQ**) the DFG motif residue is absent. Panel E (**1B38**) is type I but shows a DFG-in conformation with an αC-out state. In panel F (**3RHK**), the inhibitor binds in a type I mode despite both the DFG and αC-helix adopting out conformations. Panels G (**5TCO**) and H (**3B2W**) display DFG-in and αC-in states, contradicting the expected back-pocket binding of type II inhibitors. Together, these cases illustrate structural exceptions where rigid rule-based classification schemes fail.

### 2.5 Supervised Machine Learning Classification

The machine learning classifier was constructed by combining an 85-dimensional continuous distance feature vector, describing residue-level pocket geometry across conserved kinase structural motifs, with three categorical conformational state annotations (DFG conformation, αC-helix state, salt bridge status). Continuous distance features were preprocessed using median imputation to handle missing values arising from unresolved residues in individual structures and Categorical features were imputed using the most frequent category using scikit-learn[41]. The rule-based labeled dataset (6,485) was screened for exact duplicate rows and label conflicts prior to training. The target variable comprised seven classes: Type I, I½, II, III (Top, Back, Front) and Type IV.

A tiered oversampling strategy was applied on the training partition only, majority classes (Type I and IV) were left unaltered but the others were oversampled, capped at 500 samples per class, using the Synthetic Minority Oversampling Technique (SMOTE)[42]. Tomek Links undersampling was subsequently applied to remove noisy majority-class boundary samples[43]. The labeled dataset was partitioned into training and test (80:20) sets using stratified random sampling. Seven candidate classifiers Random Forest[44], Extra Trees[45], Gradient Boosting[46], Logistic Regression, Support Vector Machine[47], k-Nearest Neighbors[48], and XGBoost[49] were compared on the SMOTE balanced training data, the Extra Trees classifier achieved the best cross-validated performance and was selected as the final model architecture. Hyperparameters for the Extra Trees classifier used comprised 200 trees, unrestricted tree depth, a minimum split size of 2, and √(total features) considered per split, with balanced class weighting.

The tuned model was evaluated on the untouched, non synthetic held out test set, achieving an overall accuracy of 0.992, weighted F1-score of 0.992, and macro-averaged F1-score of 0.941. Per-class performance was near perfect, with only little classification uncertainty occurring among the structurally adjacent Type III subtypes (Top, Back, Front) and at the Type I and Type I½ boundary, consistent with the mechanistic ambiguity of these binding modes. Feature importance derived from the tuned ensemble identified DFG conformation state, hinge-region distances, and salt-bridge status as the most discriminative descriptors of binding mode, in agreement with established structural determinants of kinase-inhibitor recognition. The validated model was applied to predict binding modes for the 5,460 previously unclassified kinase-ligand complexes, extending classification coverage to the full curated dataset of 11,945 kinase-ligand pairs. The majority of these predictions were made with high confidence.

### 2.6 Phylogenetic Integration and Druggability Transfer

To investigate the relationship between kinase evolutionary history and inhibitor binding mode accessibility, the ML derived binding mode classifications were integrated with a reference kinome phylogeny. This integration enabled a kinome-wide visualization of inhibitor type distributions mapped onto the kinase phylogenetic tree, and phylogenetic druggability transfer- the hypothesis that evolutionary proximity between kinases predicts shared pocket accessibility and, by extension, shared susceptibility to specific inhibitor binding modes.

Kinase phylogenetic relationships were represented using the canonical human kinome reference tree obtained from KinHub[50], derived from the landmark kinome classification of Manning et al., which defines the eight major kinase groups: AGC, CAMK, CK1, CMGC, RGC, STE, TK, and TKL, together with a group of atypical and other kinases (OTHER). This provides a standardized evolutionary scaffold against which structural and pharmacological data can be systematically mapped.

Each kinase entry in the classified dataset was assigned to its corresponding node in the kinome tree via its UniProt accession identifier. Since the reference kinome tree is anchored on human kinases, non-human kinase structures present in the dataset were assigned to their human ortholog. This approach is justified by the high conservation of kinase domain architecture across eukaryotes, particularly at the catalytic cleft residues constituting the 85-residue binding site framework. Kinase entries that could not be unambiguously resolved to a node in the reference tree were excluded from phylogenetic mapping analyses but retained in the broader structural atlas.

To visualize the global distribution of inhibitor binding modes across the kinome, binding mode classification data were mapped onto the kinome tree using the CORAL web server[51]. To provide granular phylogenetic resolution within each kinase group, eight separate circular kinome tree visualizations were generated, one for each of the eight canonical kinase groups (AGC, CAMK, CK1, CMGC, RGC, STE, TK, TKL). In each group-level tree, kinase nodes were displayed as leaves of the tree, with outer annotation rings encoding the relative abundance of each inhibitor type observed for that kinase. This visualization framework enables qualitative assessment of whether specific inhibitor binding modes cluster within particular kinase families or clades or whether they are distributed in a manner uncorrelated with evolutionary relatedness. Phylogenetic druggability signal was quantified by computing binding mode co-occurrence rates between kinase pairs across equal count phylogenetic distance bins, compared against a permutation null distribution (1,000 permutations), enrichment was quantified as a permutation Z-score. This approach operationalizes the evolutionary target proximity framework of Chakraborti et al. [29] in the context of allosteric pocket accessibility prediction.

## 3. Results and Discussion

### Conformational Landscape of the Kinase Atlas

Systematic curation yielded an atlas of 12,641 kinase domain structures spanning 477 UniProt kinase domains from 467 UniProt accessions across diverse eukaryotic organisms a comprehensive organism-spanning kinase structural atlas assembled for systematic inhibitor classification, substantially exceeding the scope of KLIFS[24], [52] with only human and mouse kinase and the reviewed entries of KinCore dataset as well [26]. Application of a ligand-presence filter, with exclusion of solvent molecules and crystallographic additives, identified 10,739 structures containing qualifying small-molecule ligands. After decomposing multi-ligand entries into independent protein-ligand pair records, the final analytical dataset comprised 11,945 unique kinase-ligand complex records, each defined by a specific PDB entry, chain, and ligand instance.

Conformational annotation revealed the expected kinome-wide dominance of DFG-in (∼90%), DFG-out in (∼8.0%), and DFG-out-like intermediate in (∼02.0%) is in quantitative agreement with Ung et al.’s Kinformation analysis (88.5% DFG-in, 9.0% DFG-out, 2.5% intermediate)[30], independently validating our geometric descriptor thresholds. The known PDB bias toward DFG-in conformations[53] has important implications: inactive DFG-out conformations represent only a fraction of available structural templates, reinforcing the value of ML-based pocket accessibility inference and AI-guided conformational sampling for understudied kinases. Consistent with the prevalence of DFG-in structures, αC-helix-in configurations were observed in the majority of structures ∼94.5%, with αC-out conformations accounting for ∼5.5%. The catalytic salt bridge was present in ∼64% of all structures overall, with markedly higher rates in CK1 (91.9%), AGC (87.7%), CAMK (79.5%), and CMGC (78.4%) kinases, consistent with the hydrophobic spine model[16] predicting predominantly active conformations for these groups and substantially lower in atypical kinases (16.8%), which employ non-canonical regulatory architectures[54]. These group-level conformational differences directly predict and are validated by the group-level inhibitor binding mode distribution patterns observed downstream (Table S6).

**Figure 5.**
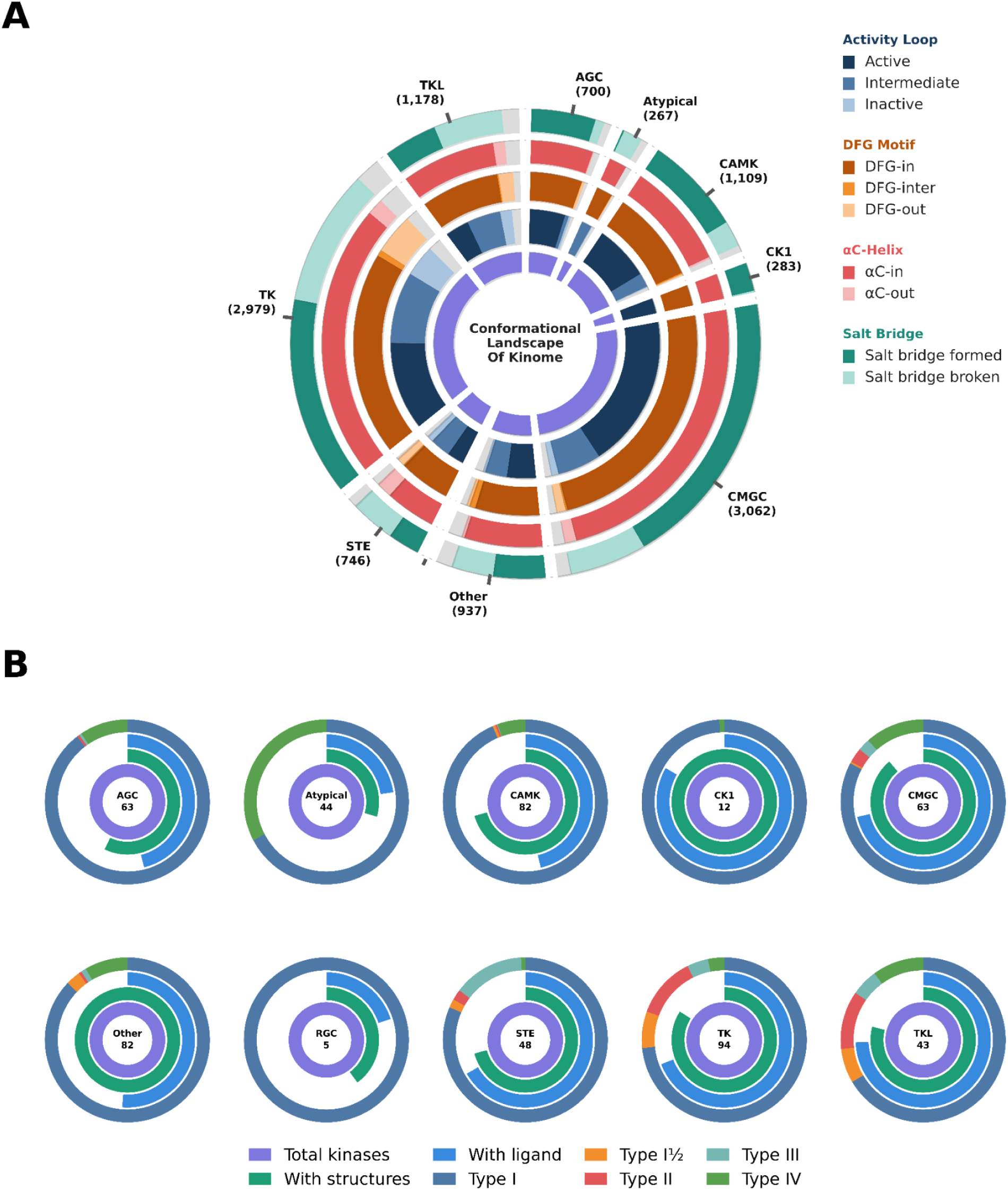
Kinome-wide conformational and pharmacological landscape of all structural atlas. (A) Concentric ring chart showing, from inner to outer rings: Total PDB structures, activation state (Active / Intermediate / Inactive), DFG conformation (DFG-in / DFG-out-like / DFG-out), αC-helix position (in / out) and salt-bridge status for all kinase groups (n shown per group). (B) Per-group donut charts displaying, from inner to outer rings: Total known kinases, kinases with resolved structures, kinases with bound ligands, and the relative proportions of each inhibitor binding-mode class from the structural data (Type I-IV, colour-coded as in legend). Numbers inside donuts indicate unique kinases in each group.

### Machine Learning Classification of Kinase-Ligand Binding Modes

The rule-based framework generated a high-confidence training set Type I (∼71.9%), Type IV (14.8%), Type II (6.0%), Type III (4.8%), Type I½ (2.6%), consistent with the broader kinase inhibitor literature in which ATP-competitive agents dominate approved drugs while allosteric classes remain underrepresented relative to their selectivity potential[8], [11], [55]. The optimized Extra Trees classifier achieved an overall test accuracy of 99.15% with weighted F1 = 0.992, macro-F1 = 0.941, indicating strong and balanced predictive performance. Ten-fold cross-validation on the balanced training data confirmed this performance (accuracy 0.996, macro-F1 0.995), and learning curves showed a training-validation macro-F1 gap of only 0.005, indicating no overfitting despite the use of synthetic minority samples. After refitting on the full labelled set, the model was applied to the 5,460 previously unresolved complexes, extending full binding-mode coverage to all 11,945 kinase-ligand pairs. Predicted labels for this subset followed a distribution broadly consistent with the labelled set (Type I ∼85.7%, Type IV ∼5.5%, Type II ∼4.2%, Type I½ ∼3.3%, Type III sub-pockets ∼1.2% combined), and prediction confidence was high overall with 4,083 complexes (74.8%) were classified at >90% probability and a further 1,084 (19.9%) at 70–90%, with only 38 complexes (0.7%) having low prediction confidence.

feature importance identified DFG conformational state (DFG-in and DFG-out indicators) as the strongest discriminative signal, followed closely by hinge-region distances, salt-bridge status, and αC-helix position. This ranking is mechanistically coherent, hinge hydrogen bonding defines the boundary between ATP-competitive (Types I, I½, II) and ATP-non-competitive (Types III, IV) binding modes, DFG-loop conformation directly encodes the classical Type I versus Type II distinction and the pocket rearrangement that Type II ligands exploit, and αC-helix position is the structural feature that separates Type I½ from Type I and typically accompanies DFG-out pocket formation in Type II binding. The convergence of the model’s data-driven feature ranking with established structural pharmacology supports the conclusion that the classifier is learning mechanistically grounded signals rather than spurious correlations.

Mapping ML-classified binding modes onto the Manning kinome reference tree[5] revealed group-level patterns of inhibitor accessibility reflecting the combined influence of conserved structural architecture, clade-specific conformational plasticity, and evolutionary history. These patterns carry direct mechanistic and pharmacological implications.

**Type I** inhibitors are broadly distributed across all eleven kinase groups (CMGC 2,305; TK 2,125; CAMK 834), consistent with universal ATP-binding hinge conservation and the well-documented polypharmacology of ATP-competitive agents across structurally diverse kinase targets[56]. The structural basis of this universality is the conserved hinge backbone hydrogen bond acceptor/donor pattern, which is maintained even in kinases with highly divergent sequences flanking the ATP site.

**Type I½** inhibitors show a strikingly restricted distribution: 60.9% of 338 structures map exclusively to TK, 23.3% to TKL. This concentration reflects the unusually high αC-helix mobility in receptor tyrosine kinases documented by Shan et al. through MD simulation,[57] a clade-specific conformational property mechanistically linked to the hydrophobic spine architecture of TK kinases where spine-stabilizing residues diverged at the TK/TKL branch point[19]. The near-complete absence of Type I½ structures in AGC, CAMK, and CK1 kinases where the αC-helix is stabilized by a rigid hydrophobic spine[16] is consistent with that αC-helix-mediated pocket accessibility being a phylogenetically heritable property. This is a pharmacologically actionable finding: it supports deprioritizing AGC and CAMK kinase targets for Type I½ design strategies, saving significant lead optimization effort.

**Type II** inhibitors are concentrated in TK (348, 58.6%) and TKL (135, 22.7%), with notable CMGC representation (84, 14.1%) dominated by p38α. The near-complete absence in AGC and CAMK kinases is consistent with structural evidence that the DFG-out pocket in AGC kinases is sterically obstructed by the intact hydrophobic spine[35], [54]. This supports the pharmacological inference that AGC kinases involved in cancer-driving pathways (AKT, RSK, PKA) are inherently resistant to DFG-out targeting strategies, and suggests allosteric approaches via the PIF-pocket, PH-domain interface, or other Type IV mechanisms may be more productive design vectors for this group. The phylogenetic clustering of Type II accessibility within TK and TKL clades validates that DFG-out propensity is a clade-level structural property determined by evolutionary conservation of the x-DFG position and hydrophobic spine residues, as first characterized by Ung et al.[30]

**Type III** inhibitors exhibit the most restricted distribution- 33.0% in TK, 27.2% in STE, 19.2% in CMGC and 16.8% in TKL. Within each group, Type III structures cluster tightly within specific clades rather than distributing uniformly. The STE enrichment is almost entirely attributable to MEK1/2, whose unique allosteric cavity formed partly by a MEK-specific insert between β and αC structural elements absent in other kinases[58], [59], represents one of the most clade-restricted allosteric binding sites in the kinome. The joint absence of both Type II and Type III structures in most AGC and CAMK kinases is consistent with the rigid hydrophobic spine and compact activation loop geometry of these groups, which likely obstruct both the DFG-out back pocket and the adjacent allosteric space. This pattern points toward Type IV remote allosteric strategies as a more promising avenue for drug discovery in these kinase families.

**Type IV** inhibitors show the broadest distribution: CMGC 469 (43.1%), TKL 249 (22.8%), AGC 64 (5.8%). The appearance of Type IV structures in kinases lacking Types II and III is consistent with remote allosteric sites at regulatory interfaces being a viable pharmacological strategy for these kinase families, in line with predictions from the Pocket-Phylogeny-Function axis framework.

**Figure 6.**
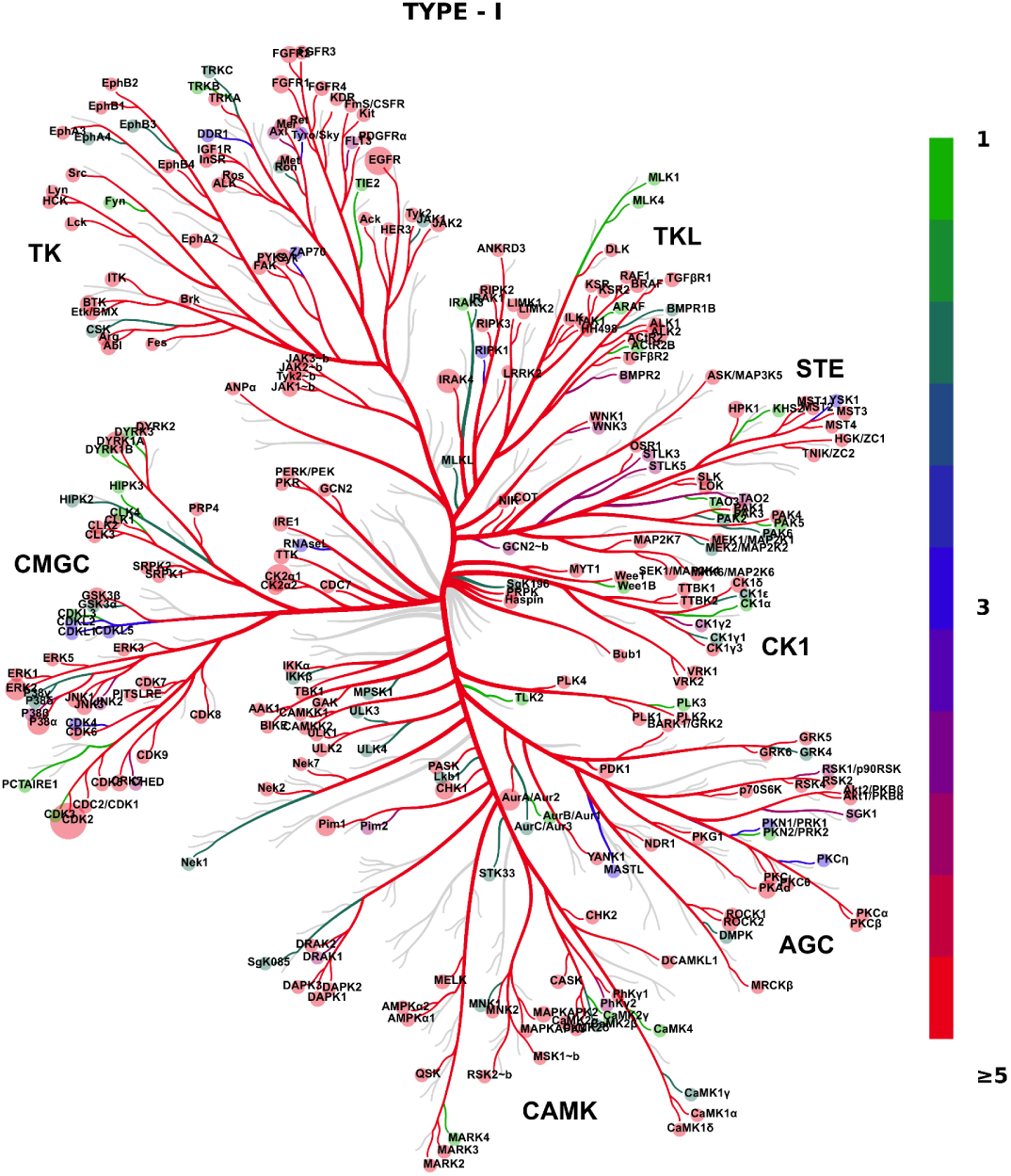
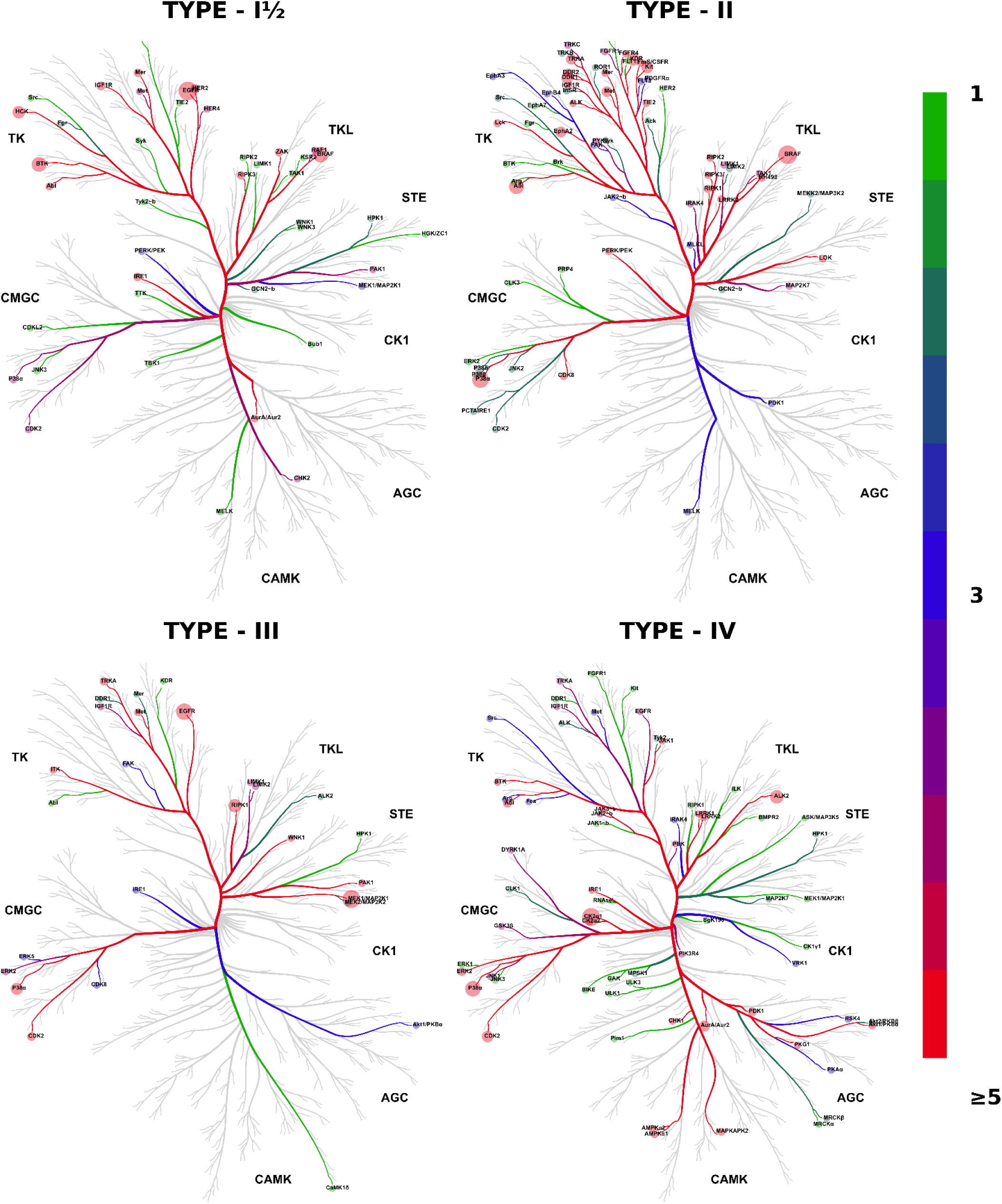
Kinome wide phylogenetic distribution of inhibitor binding modes reveals clade-stratified pharmacological accessibility. Five Manning kinome trees rendered via CORAL, one per inhibitor class (Type I, I½, II, III and IV). Node colour is proportional to the number of co-crystal structures of each inhibitor class observed for that kinase (shown on scale). kinase with no instance of inhibitor are in grey.

### Pharmacological Validation of Allosteric Pocket Subclasses

To demonstrate that the proposed allosteric pocket subclasses represent biologically meaningful pharmacologically validated binding sites, approved or clinically advanced inhibitor-bound structures with experimentally established binding modes were mapped to each proposed subclass. The systematic co-occurrence of structurally diverse approved agents within these predicted subclasses provides strong evidence that the spatial partitioning framework captures genuine pharmacological topography rather than geometric artifacts.

#### Type III-Front

The Type III-Front subclass captures ligands occupying the hydrophobic allosteric cavity adjacent to the ATP site while extending across the activation loop toward the front cleft. Nine FDA-approved or clinically advanced MEK inhibitors with available co-crystal structures were assigned exclusively to this subclass: trametinib (7JUX), selumetinib (4U7Z, 7M0T), binimetinib (7M0U), cobimetinib (4AN2), mirdametinib (3EQG), pimasertib (7M0W), refametinib (3E8N), and avutometinib (3WIG). All make critical contacts with the activation loop, αC-helix and hydrophobic spine residues, stabilizing the inactive MEK conformation, as characterized by Ohren et al. and Fischmann et al.[58], [59], The exclusive enrichment of nine chemically diverse approved inhibitors within a single subclass provides unambiguous pharmacological validation.

#### Type III-Back

The Type III-Back subclass consists of ligands positioned entirely behind the activation loop within the deep hydrophobic back cleft. This subclass displayed binding geometries distinct from Type III-Front inhibitors, supporting the existence of a separate spatially constrained allosteric subpocket. Although there are research-stage candidates but this region currently lacks approved drugs or clinical candidates, A representative structure is PDB: 8FP0 ligand ID 7TW.

#### Type III-Top

The Type III-Top subclass defines ligands positioned above the activation loop,the EGFR inhibitor naquotinib, PDB: 7LG8 exhibits the Top binding geometry in which the ligand extends upward across the activation loop. The successful identification of this distinct binding arrangement further supports the structural granularity of the Type III subclassification scheme.

#### Type IV-A: Ibrutinib Binding Site

The Type IV-A subclass corresponds to the distal non-canonical binding site occupied by Bruton’s tyrosine kinase (BTK) inhibitors such as ibrutinib. The PDB: 6YG2 structure confirmed localization within the experimentally validated BTK regulatory pocket distinct from the canonical ATP-binding site, supporting the specificity of the Type IV-A definition.

#### Type IV-B: αD-Pocket

The Type IV-B subclass represents ligands occupying the regulatory αD-pocket adjacent to the αD-helix. The identified structure PDB: 8QBU demonstrated a discrete distal binding cavity structurally separated from both ATP and activation-loop-associated allosteric regions, supporting classification as an independent allosteric pocket subclass.

#### Type IV-C: PIF-Pocket

The Type IV-C subclass corresponds to the conserved PIF-pocket (PDK1-interacting fragment pocket), a hydrophobic regulatory groove involved in kinase activation and substrate recognition. (5LVO) The assigned ligand occupied the canonical hydrophobic motif-binding cleft characteristic of PIF-pocket interactions, confirming the structural validity of this subclass.

#### Type IV-D: Regulatory Myristate Binding Site

The Type IV-D subclass represents ligands binding the regulatory myristoyl pocket located within the kinase C-lobe, particularly characteristic of ABL-family kinases. Clinically relevant inhibitors assigned to this subclass included drugs like Bosutinib (6FDY), and Ponatinib (3ZOS). These inhibitors occupy the experimentally established myristate regulatory cavity associated with conformational control of ABL kinases. Their consistent subclass assignment supports the ability of the framework to identify distal regulatory lipid-binding pockets.

#### Type IV-E: PH-Domain Allosteric Pocket

The Type IV-E subclass defines an allosteric interface formed between the kinase catalytic domain and the pleckstrin homology (PH) domain. Miransertib (5KCV) is a well-characterized allosteric AKT inhibitor that stabilizes the inactive PH-in conformation by occupying the interdomain regulatory pocket.

#### Type IV-F: Lower Myristoyl Pocket

The Type IV-F subclasses correspond to ligands targeting the bottom part of regulatory myristoyl-associated pockets within ABL-family kinases. Asciminib (8SSN), the first FDA-approved specifically allosteric ABL1 inhibitor, binds in this myristoyl pocket and stabilizes the inactive kinase conformation independently of ATP-site occupancy.

Most subclasses capture a distinct allosteric binding mode associated with approved or clinically advanced kinase inhibitors, thereby supporting the biological validity of the proposed pocket taxonomy.

**Figure 7.**
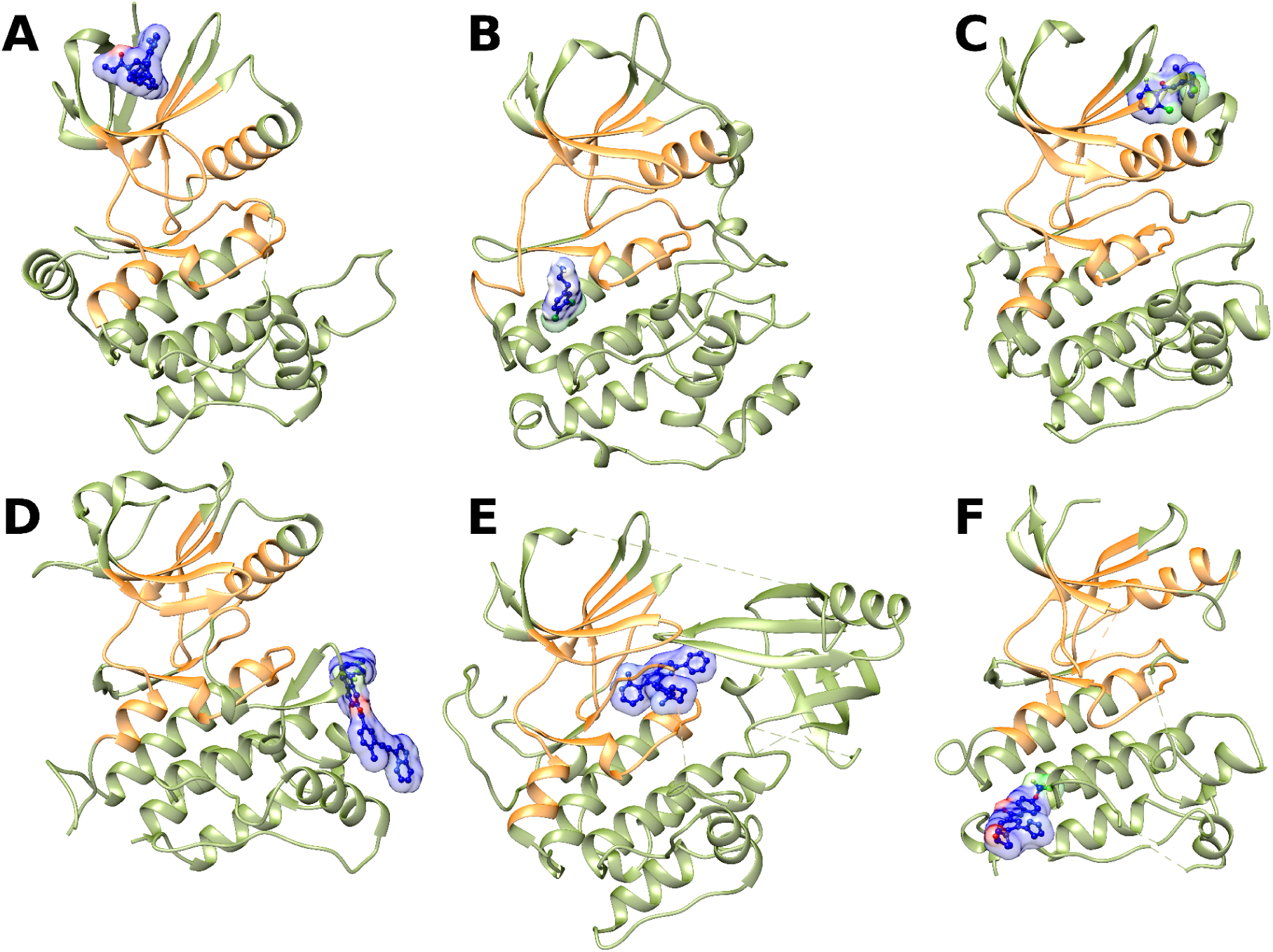
Representative co-crystal structures illustrating Type IV allosteric pocket subclasses. (A) Type IV-A: Ibrutinib Binding Site (PDB: 6YG2), (B) Type IV-B: αD-Pocket (PDB: 8QBU), (C) Type IV-C: PIF-Pocket: (PDB: 5LVO), (D) Type IV-D: Regulatory Myristate Binding Site (PDB: 3ZOS), (E) Type IV-E: PH-Domain Allosteric Pocket (PDB: 5KCV), (F) Type IV-F: Lower Myristoyl Pocket (PDB: 8SSN). Inhibitors shown as blue surfaces and kinase catalytic domain in orange.

### The Pocket-Phylogeny-Function (PPF) Axis

The convergent evidence from kinome-wide binding mode mapping, phylogenetic druggability analysis, and structural biology establishes a unified three-way relationship between pocket geometry, phylogenetic relatedness, and functional conformational behavior the Pocket-Phylogeny-Function (PPF) axis. This framework generates pharmacological inferences that collectively reposition the kinome phylogeny as a map of pharmacological opportunity.

#### 1: Phylogenetically related kinases share pocket geometry AND functional conformational plasticity

The phylogenetic druggability signal is not merely a reflection of shared primary sequence similarity but also reflects shared conformational dynamics. A kinase evolutionarily related to one with a characterized DFG-out pocket is predicted not only to have structurally similar residues lining that pocket, but to have comparable propensity to sample the DFG-out conformation during its conformational ensemble. Phylogenetic classification therefore predicts both pocket existence (structural) and pocket accessibility (functional/dynamic), the two distinct prerequisites for productive inhibitor binding. The direct experimental support for this inference comes from Shan et al.’s MD simulations[57] showing that TK-group kinases sharing phylogenetic proximity sample similar αC-helix displacement trajectories, and from the Kinformation analysis[30] demonstrating that DFG position identity is both phylogenetically conserved and predictive of DFG-out propensity.

#### 2: Functional pathway overlap between phylogenetically related kinases amplifies the pharmacological impact of shared allosteric pockets

Phylogenetically related kinases tend to operate within the same or connected signaling pathways, phosphorylate overlapping substrate motifs, and exhibit partial functional redundancy.[19], [20] This means that an allosteric inhibitor series developed for one kinase within a clade is likely applicable to phylogenetically proximate neighbors operating in the same signaling context. The MAPK cascade provides the clearest example: MEK1 and MEK2, which share the Type III-Front allosteric pocket, also share ERK-directed signaling function and oncogenic redundancy[60]. An allosteric inhibitor targeting the shared MEK pocket simultaneously disrupts functional redundancy at the pathway level, offering both pharmacological selectivity (through kinase-specific pocket geometry) and therapeutic robustness (through dual kinase suppression). This principle extends to other kinase clades, phylogenetically proximate TK-group kinases sharing DFG-out accessibility and participating in the same receptor tyrosine kinase signaling axis are prime candidates for dual Type II targeting strategies that exploit both structural and functional conservation, preventing the acquired resistance driven by compensatory pathway activation through related kinases.

#### 3: Phylogenetic pocket conservation defines a selectivity hierarchy across inhibitor classes

Type I inhibitor selectivity is inherently limited because the hinge interface is conserved across the entire kinome. Type II selectivity is achievable within but not across phylogenetic groups, as DFG-out pocket geometry is clade-conserved but diverges between distant groups. Type III selectivity is the highest achievable through the KLIFS-defined binding modes, as the adjacent allosteric pocket exists only within narrow evolutionary lineages. Type IV selectivity is inherently maximal, as remote allosteric sites at regulatory interfaces are often unique to individual kinases or very small subfamilies. This hierarchy of selectivity matches the phylogenetic specificity observed in our permutation Z-score analysis providing quantitative, data-driven validation of the intuitive pharmacological principle that allosteric targeting achieves superior selectivity through less conserved binding sites. This framework directly informs drug design strategy: when designing against a kinase in the TK group, Type II or III allosteric approaches will achieve substantially better selectivity than Type I, and the phylogenetic context predicts which approach is structurally accessible.

#### 4: Dark kinases can be systematically prioritized for allosteric programs based on phylogenetic context

Of the 518 human kinases, approximately 153 (31%) have no experimentally determined crystal structure[53]. For these dark kinases[61], conventional structure-based drug design is infeasible. However, the PPF axis converts this challenge into an opportunity: a dark kinase’s phylogenetic position, combined with binding mode accessibility characterized for its experimentally solved neighbors, provides a probabilistic prediction of which allosteric pocket classes are accessible. Kinases within TK or TKL groups near characterized Type II-rich kinases can be confidently prioritized for DFG-out co-crystallization in relevant inactive conformations. Kinases within MEK-proximate STE group clades can be prioritized for Type III allosteric screening. This systematizes what has historically been unsystematic expert judgment, converting collective structural knowledge into quantitative druggability predictions. It also complements AI-based conformational sampling: while AlphaFold2 models kinases predominantly in DFG-in conformations[53]; limiting their utility for allosteric pocket identification the PPF framework guides targeted MD or AF3-based conformational sampling toward the inactive conformations phylogenetically predicted to be accessible, maximizing the information yield from computational structure prediction.

### Druggability Prioritization of Uncharacterized Kinases

Application of the phylogenetic druggability correlation across all kinases lacking co-crystal structures can generate a kinome-wide prioritization map for each inhibitor class. Kinases predicted to possess accessible Type II or Type III binding modes based on the evolutionary proximity of structurally characterised neighbours represent the highest priority targets for structure based allosteric inhibitor programs in the absence of direct structural data. This framework converts the collective structural knowledge embedded in the kinase co-crystal structure atlas into actionable predictions for under explored kinases, directly operationalizing the phylogenetic druggability transfer principle as a prospective drug discovery tool.

### Limitations and Future Directions

A few limitations of the present work merit consideration. The dataset is inherently biased toward kinases of high pharmaceutical interest, with TK and CMGC kinases accounting for a disproportionate fraction of available structures. This bias means that the phylogenetic druggability predictions are most reliable for kinases in well-represented clades and should be interpreted with greater caution for structurally understudied groups. Future work could include the development of a quantitative phylogenetic druggability scoring framework that systematically ranks kinase targets based on the evolutionary conservation and predicted accessibility of specific binding site architectures across inhibitor classes. A further limitation of this work is that the structural atlas is built entirely from publicly available PDB structures from RCSB, and is therefore blind to the limited amount of proprietary structural data held within pharmaceutical and biotechnology organisations. Industry datasets routinely contain unpublished kinase structures, including novel allosteric complexes. The phylogenetic druggability framework developed here is designed to accommodate such data. Incorporating even a modest number of proprietary allosteric structures for any understudied kinase clade is expected to substantially improve druggability predictions for the surrounding evolutionary neighbourhood.

## 4. Conclusions

We present a machine learning guided structural atlas of 11,945 kinase-inhibitor complexes that extends binding-mode classification coverage across the human kinome, resolving seven mechanistically defined classes, including three novel Type III allosteric subclasses and six Type IV remote allosteric subclasses with >99% classifier accuracy. By integrating these classifications with Manning kinome phylogeny, we show that evolutionary proximity is a statistically significant predictor of shared allosteric pocket accessibility. Together with the Pocket-Phylogeny-Function axis, this framework repositions kinome phylogeny from a taxonomic reference into an actionable prioritization tool for the roughly 153 human kinases lacking solved structures, it offers a testable, quantitative basis for selecting which allosteric strategy (Type II, III, or IV) is most likely to be structurally accessible, rather than relying on case by case expert judgment. The framework is extensible: incorporation of proprietary industry structures, prospective AlphaFold3 or MD-guided conformational sampling of predicted accessible pockets, and expansion of the phylogenetic model beyond the human kinome to disease relevant pathogen or plant kinomes are natural next steps. More broadly, this work argues that selectivity in kinase drug discovery is not solely a chemistry problem to be solved compound by compound, but one that can be substantially de-risked upstream by evolutionary reasoning about pocket accessibility.

