## Supplemental Table 1 for "Machine Learning-Guided Classification of Druggable Pockets and Phylogenetic Druggability Transfer Across the Human Kinome"

| State | DFG Conformation | αC-Helix | Salt Bridge |
|---|---|---|---|
| **Active** | DFG-in | In (≤13.15 Å) | Present (<4.0 Å) |
| **Intermediate** | DFG-in or DFG-out-like | In (≤13.15 Å) | Absent bridge |
| | DFG-out-like | In (≤13.15 Å) | Present (<4.0 Å) |
| | DFG-in | Out (>13.15 Å) | - |
| **Inactive** | DFG-out | - | - |
| | DFG-out-like | Out (>13.15 Å) | Absent bridge |

***Table S1.*** *decision logic for kinase activation state assignment based on combined conformational descriptors.*

### Machine Learning Classification of Kinase Inhibitor Binding Modes

#### S1. Dataset Composition and Class Definitions.

Seven inhibitor binding mode categories were defined based on the established kinase inhibitor classification framework. The final labelled dataset consisted of 6,481 structures with confirmed binding mode annotations from rules and manual check. An additional 5,462 structures without assigned categories were retained as an independent set and subsequently classified using the final trained model.

| Inhibitor Class | n (Labelled) | % of Dataset |
|---|---|---|
| Type 1 | 4,663 | 71.90% |
| Type 4 | 957 | 14.76% |
| Type 2 | 388 | 5.98% |
| Type 1.5 | 167 | 2.58% |
| Type 3 Back | 130 | 2.00% |
| Type 3 Top | 100 | 1.54% |
| Type 3 Front | 80 | 1.23% |

***Table S2.*** *Dataset class composition. Maximum class imbalance ratio: 58:1 (Type 1 vs. Type 3 Front).*

#### S2. Class Imbalance Handling

A tiered SMOTETomek resampling strategy was applied exclusively to the training partition. Complete balancing of all classes to match the majority class size (3,730 samples) was intentionally avoided, as this would have generated datasets composed predominantly of synthetic observations. For example, the Type 3 Front class contained only 64 original training samples and would have required expansion to 3,730 samples, resulting in more than 98% synthetic data and increasing the risk of overfitting to SMOTE-generated interpolation artefacts. Instead, tiered resampling targets were defined as follows

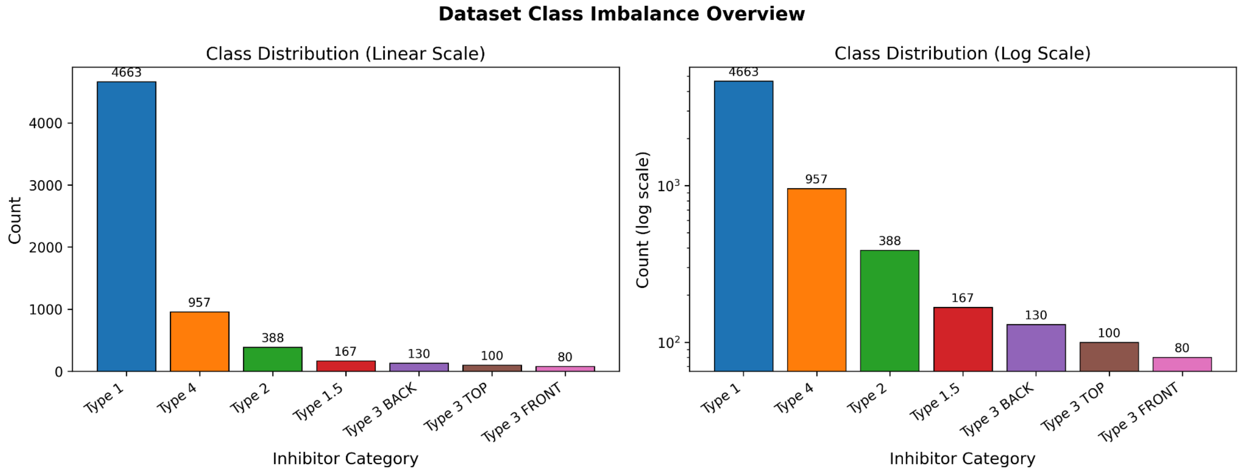

***Figure S1***. *Overview of class imbalance in the labelled kinase inhibitor dataset (n = 6,481).Class distributions are shown on linear (left) and logarithmic (right) scales.*

| Class | Original | train | Target after SMOTE | Multiplier | Synthetic % |
|---|---|---|---|---|---|
| Type 3 Front | 80 | 64 | 500 | 7.8x | 87% |
| Type 3 Top | 100 | 80 | 500 | 6.2x | 84% |
| Type 3 Back | 130 | 104 | 500 | 4.8x | 79% |
| Type 1.5 | 167 | 134 | 500 | 3.7x | 73% |
| Type 2 | 388 | 310 | 930 | 3.0x | 67% |
| Type 4 | 966 | 766 | 766 | 1.0x | 0% |
| Type 1 | 4,651 | 3,730 | 3,730 | 1.0x | 0% |

***Table S3***. *Tiered SMOTETomek resampling strategy applied to training data. Tomek Links were applied after SMOTE to remove borderline majority-class samples near decision boundaries. class_weight='balanced' was additionally applied within the Extra Trees classifier as a secondary guard.*

### S3. Test Set Performance Per-Class Report

| Class | Precision | Recall | F1-Score | Support (n) |
|---|---|---|---|---|

| | | | | |
|---|---|---|---|---|
| Type 1 | 1 | 1 | 1 | 933 |
| Type 1.5 | 1 | 0.88 | 0.94 | 33 |
| Type 2 | 1 | 1 | 1 | 78 |
| Type 3 Back | 0.96 | 0.88 | 0.92 | 26 |
| Type 3 Front | 0.88 | 0.88 | 0.88 | 16 |
| Type 3 Top | 0.82 | 0.90 | 0.86 | 20 |
| Type 4 | 1 | 1 | 1 | 191 |
| Accuracy | | | 0.99 | 1,297 |
| Macro avg | 0.95 | 0.93 | 0.94 | 1,297 |
| Weighted avg | 0.99 | 0.99 | 0.99 | 1,297 |

***Table S4***. *Per-class classification performance on the held-out test set (n = 1,297 i.e. 20% stratified split). Support denotes the number of real samples in each test class.*

### S4. Performance Metrics

| Metric | Value |
|---|---|
| Test Accuracy | 0.9915 |
| F1-Score (Weighted) | 0.9915 |
| F1-Score (Macro) | 0.9408 |
| Matthews Correlation Coeff. | 0.9813 |
| Final Train F1-Macro | 1 |
| Final Val F1-Macro (10-fold) | 0.9952 |
| Train–Val Gap | 0.0048 |

***Table S5.*** *Summary performance metrics for the Extra Trees classifier on the held-out test set.*

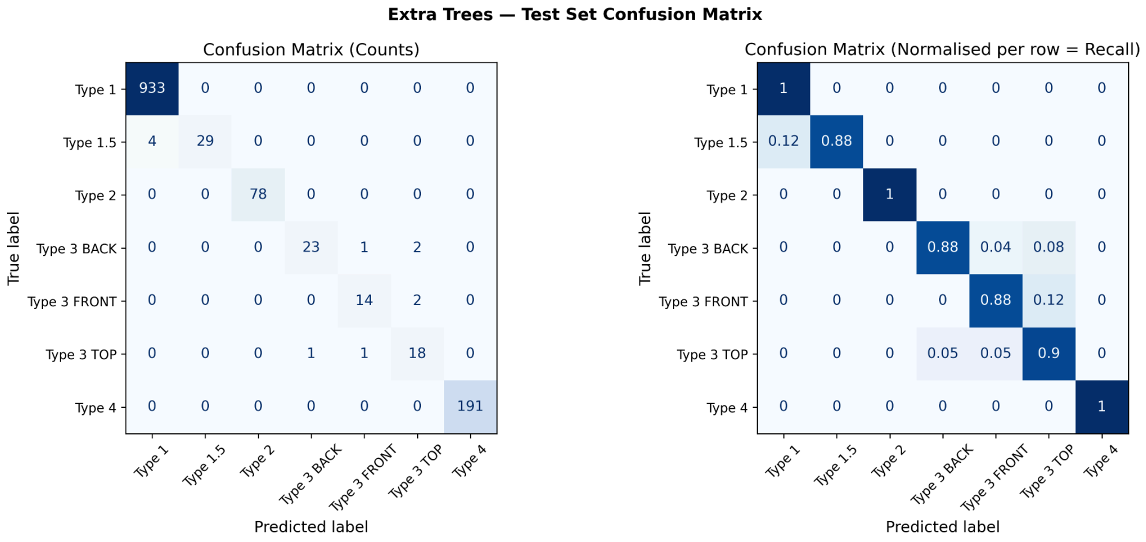

***Figure S2***. *Confusion matrix for the Extra Trees classifier on the held-out test set (n = 1,297). Left: raw prediction counts. Right: row-normalised matrix. Type 1, Type 2, and Type 4 achieve perfect or near-perfect recall. Residual misclassification is confined exclusively to within the Type 3 subtype triad, which share the same allosteric pocket region and differ only in their binding face orientation.*

### S5 Feature Importances

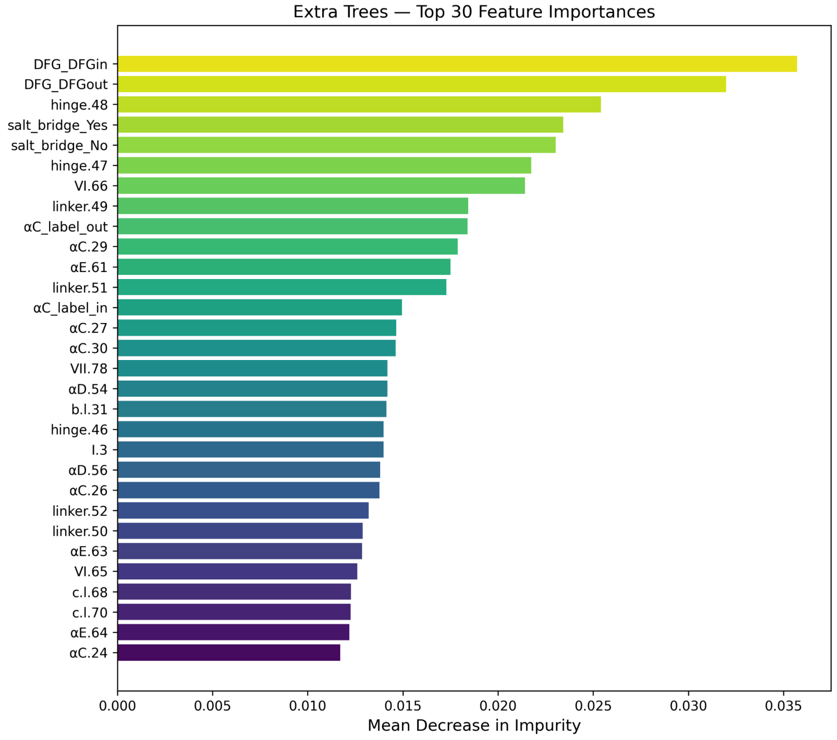

***Figure S3.*** *Top features by Mean Decrease in Impurity (MDI) from the Extra Trees classifier. DFG conformation and salt bridge and Hinge status are the most informative global features, consistent with their roles as primary determinants of kinase activation state and inhibitor binding mode.*

### S6 Learning Curves

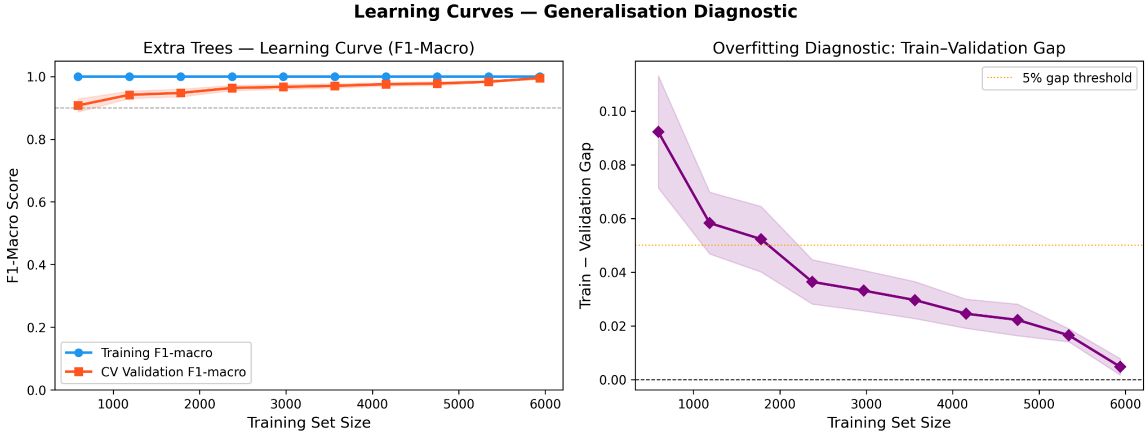

***Figure S4.*** *Learning curves for the Extra Trees classifier. Left: training (blue) and 5-fold cross-validation (orange) F1-macro scores as a function of training set size, shaded regions indicate ±1 standard deviation across folds. Right: train-validation gap (overfitting diagnostic). The gap converges to 0.007 at full training set size, well below the 5% threshold (orange dashed line), demonstrating no overfitting. The validation curve plateaus above F1 = 0.99, indicating the model has reached near-maximum performance with the current dataset.*

### S7 SHAP Beeswarm: Directional Feature Contributions

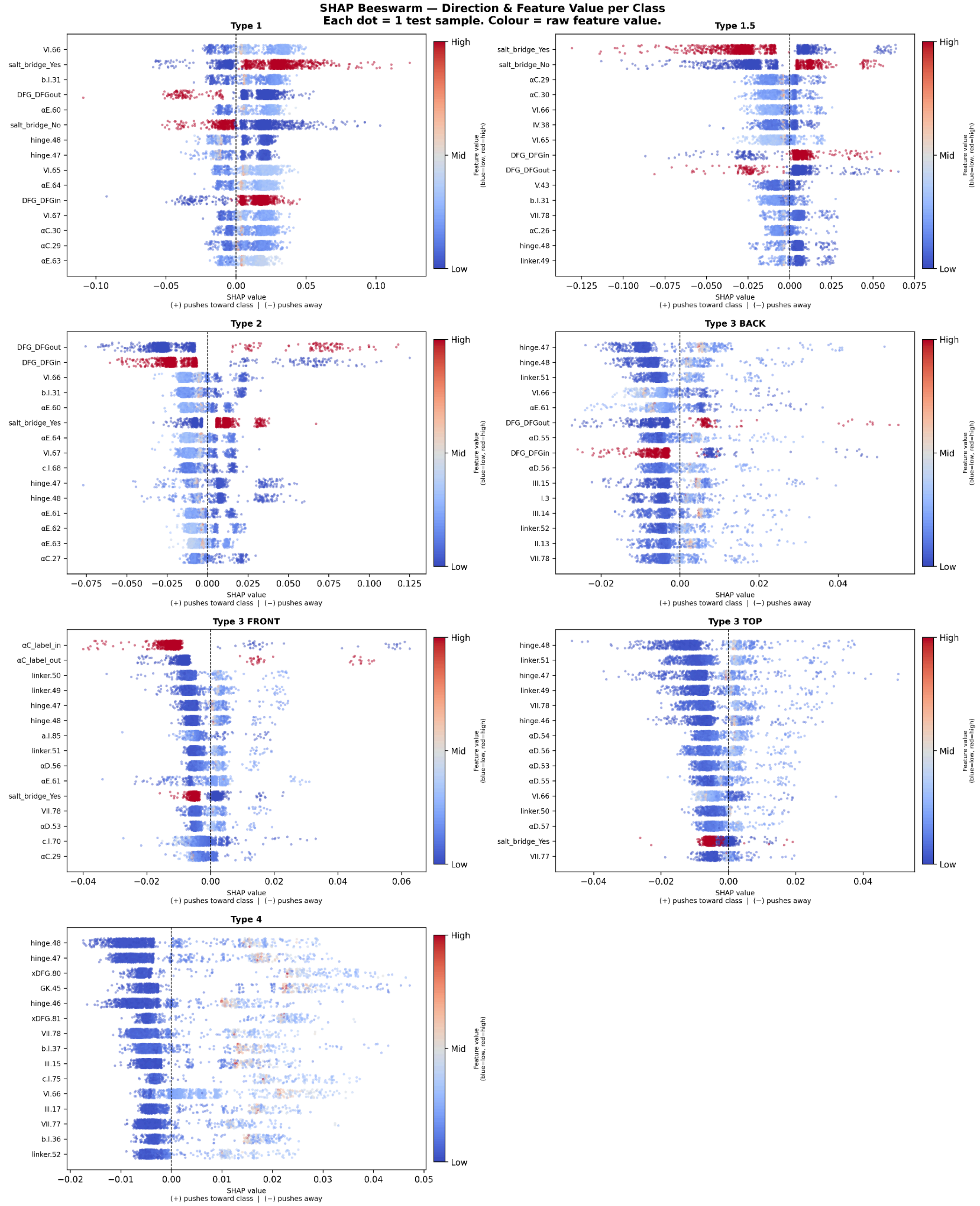

***Figure S5.*** *SHAP beeswarm plots showing signed feature contributions for each inhibitor class. Each point represents one test set structure. The x-axis indicates the SHAP value: positive values push the prediction toward the labelled class, negative values push predictions away. Point colour encodes the raw feature value (red = high , blue = low ). Features are ranked by mean absolute SHAP value.*

Key observations- Type 2 classification is driven by high DFGout values (red dots, rightward), Type 1.5 is uniquely characterised by salt bridge absence, Type 3 subtypes share hinge and αC helix features as primary drivers, with allosteric pocket contacts (low distance, blue) contributing negatively.

**S8 Distribution of kinase structures, conformational states, and inhibitor binding mode classifications across eleven kinase groups**

| | AGC | Atypical | CAMK | CK1 | CMGC | Other | RGC | STE | TK | TKL | N/A |
|---|---|---|---|---|---|---|---|---|---|---|---|
| Total Kinases domain in human | 63 | 44 | 82 | 12 | 63 | 82 | 5 | 48 | 94 | 43 | |
| Manning's human kinase | 63 | 45 | 74 | 12 | 61 | 83 | 5 | 47 | 90 | 43 | |
| Number of kinases with structures | 36 | 13 | 58 | 14 | 56 | 58 | 2 | 34 | 79 | 34 | 93 |
| Number of kinases with ligands | 29 | 10 | 38 | 10 | 45 | 42 | 1 | 32 | 65 | 32 | 73 |
| Total structures | 700 | 267 | 1109 | 283 | 3062 | 937 | 7 | 746 | 2979 | 1178 | 1373 |
| Total structures with ligands | 639 | 157 | 892 | 209 | 3048 | 887 | 6 | 713 | 2904 | 1311 | 1179 |
| Active | 540 | 30 | 821 | 251 | 2211 | 444 | 0 | 273 | 1290 | 371 | 527 |
| Inactive | 32 | 10 | 15 | 4 | 105 | 34 | 0 | 95 | 451 | 168 | 44 |
| Intermediate | 52 | 152 | 200 | 18 | 622 | 314 | 4 | 289 | 1021 | 480 | 632 |
| DFGin | 606 | 207 | 1047 | 279 | 2838 | 743 | 6 | 598 | 2298 | 896 | 1202 |
| DFGout | 30 | 10 | 10 | 4 | 105 | 34 | 0 | 94 | 451 | 164 | 44 |
| DFGout-like | 3 | 0 | 14 | 0 | 18 | 70 | 0 | 12 | 69 | 14 | 16 |
| Î±C-helix_in | 611 | 190 | 1018 | 271 | 2825 | 760 | 2 | 499 | 2600 | 906 | 1157 |
| Î±C-helix_out | 10 | 0 | 18 | 2 | 112 | 28 | 2 | 151 | 154 | 110 | 45 |
| Salt bridge-Yes | 545 | 32 | 824 | 251 | 2305 | 445 | 0 | 279 | 1640 | 457 | 549 |
| Salt bridge-No | 76 | 158 | 212 | 22 | 632 | 343 | 4 | 371 | 1114 | 559 | 653 |
| Type I | 569 | 102 | 834 | 204 | 2415 | 763 | 6 | 579 | 2125 | 787 | 960 |
| Type I½ | 0 | 0 | 5 | 0 | 10 | 25 | 0 | 13 | 206 | 79 | 8 |

| | | | | | | | | | | | |
|---|---|---|---|---|---|---|---|---|---|---|---|
| Type II | 3 | 0 | 3 | 0 | 84 | 5 | 0 | 15 | 348 | 135 | 27 |
| Type III | 3 | 0 | 1 | 0 | 70 | 9 | 0 | 99 | 120 | 61 | 14 |
| Type IV | 64 | 55 | 49 | 5 | 469 | 85 | 0 | 7 | 105 | 249 | 170 |

***Table S6***. *Distribution of kinase structures, conformational states, and inhibitor binding mode classifications across eleven kinase groups*
